# Functional and structural pathologies in skeletal muscle of a rat model of Duchenne muscular dystrophy

**DOI:** 10.1101/2025.09.11.675592

**Authors:** Young il Lee, Cora C. Hart, C. Spencer Henley-Beasley, Jeffrey S. Herr, Eli Zerpa, Elisabeth R. Barton, David W. Hammers, H. Lee Sweeney

## Abstract

**Background:** Duchenne muscular dystrophy (DMD) is a lethal pediatric degenerative muscle disease for which there is no cure. Robust preclinical models that recapitulate major clinical features of DMD are required to investigate efficacy of potential DMD therapeutics. Rat models of DMD have emerged as promising small animal models to accomplish this; however, there have been no comprehensive studies investigating the functional skeletal muscle decrements associated with the modeling of DMD in rats.

**Methods:** CRISPR/Cas9 gene editing was used to generate a dystrophin-deficient Sprague-Dawley muscular dystrophy rat (MDR). Biochemical and immunofluorescent analyses were performed to confirm loss of dystrophin in striated muscles of this rat model. *In situ* and *ex vivo* muscle function was assessed in wild-type (WT) and MDR muscles at 3, 6, and 12 months of age, followed by histopathological analyses.

**Results:** MDR muscle tissues exhibited loss of full-length dystrophin and reduced content of other dystrophin glycoprotein complex members. MDR extensor digitorum longus (EDL) muscles and diaphragms displayed pronounced and progressive muscle weakness beginning at 3 months of age, compared to WT littermates. EDLs also exhibit susceptibility to eccentric contraction-induced damage. Functional deficits in soleus muscles were less severe and were associated with a right shift in force-frequency relationship and a muscle fiber-type shift. MDR muscles display progressive histopathology including degenerative lesions, fibrosis, regenerative foci, and modest adipose deposition.

**Conclusions:** MDR is a preclinical model of DMD that exhibits many translational features of the human disease, including a large dynamic range of muscle decrements, that has high utility for the evaluation of potential therapeutics for DMD.

## INTRODUCTION

Duchenne muscular dystrophy (DMD), inherited in an X-linked recessive fashion, is a progressive muscle degenerative disease that results from mutations in the dystrophin gene (*DMD*) [1]. DMD affects ∼1:5000 male births and is the most common form of muscular dystrophy [2–4]. The subsequent lack of functional dystrophin protein at the sarcolemma of striated muscles leads to progressive muscle wasting and cardiac dysfunction [5, 6]. Individuals with DMD typically experience muscle weakness at a young age, lose ambulation in their early teens, and succumb to respiratory and/or cardiac complications. While recent translational efforts have led to multiple treatment options, including small molecules, exon skipping oligonucleotides, and adeno-associated virus (AAV)-mediated dystrophin replacement gene therapy [3, 5–7], DMD remains without a cure or treatment that completely arrests the disease progression despite its known monogenetic etiology and decades of preclinical research.

Small animal models that more accurately reflect human pathologies and progression are crucial for both understanding disease mechanisms and development of potential therapeutic strategies. The *mdx* mouse on the C57BL/10 genetic background is the most widely utilized animal model of DMD, yet, despite being a genetic homolog for DMD [1, 8], fails to replicate the severity or the pathological extent of human DMD [9–12]. This mild phenotype has resulted in the failed translation of promising preclinical data into effective clinical therapeutics [13]. The D2.*mdx* mouse (*mdx* mutation on the DBA/2J genetic background) has emerged as an improved murine model of DMD with robust muscle decrements and fibrosis [14, 15], providing a reliable platform for preclinical therapeutic testing and discovery [16–18]. However, this model still exhibits an early phase of severe degeneration and subsequent regeneration that is not a feature of the human disease [15]. While large animal models of DMD, including the golden retriever muscular dystrophy (GRMD) dog, provide an accurate progression of DMD pathology [19], the individual phenotypic variability and cost-prohibitive nature of their use limits their inclusion in rigorous preclinical evaluations. Hence, there remains a need for robust, accessible, and cost-efficient preclinical DMD models that closely mimic human pathology in the therapeutic development pipeline.

Rat models of DMD offer a promising avenue to achieve this balance of an intermediate sized species that enables both rigorous and cost-effective preclinical therapeutic development. As a result, several dystrophin-deficient rats have been developed [20–22], each able to reproduce overlapping, yet distinct, aspects of DMD in humans. This includes reduced body and muscle mass, fibrosis of striated muscle tissues, and motor tasks deficits. However, a comprehensive investigation into the skeletal muscle functional deficits associated with a rat model of DMD has not been reported, which is key to understanding the utility of this model for therapeutic testing for translational functional outcomes.

The purpose of this study is to specifically address this knowledge gap by comprehensively evaluating the skeletal muscle functional pathology associated with a CRISPR/Cas9-generated rat model of DMD, referred to as the muscular dystrophy rat (MDR). We identify significant deficits in muscle growth and force production of the respiratory and limb muscles demonstrated by the MDR model, compared to wild-type (WT) littermates, as well as the accumulation of histopathological features. These findings confirm that the MDR model of DMD exhibits a significant functional dynamic range suitable for the evaluation of potential DMD therapeutics with functional outcomes.

## RESULTS

### CRISPR/Cas9-mediated *Dmd* inactivation in Sprague-Dawley rats

A gene targeting approach was taken to alter the *Dmd* in the Sprague-Dawley rat background, preventing expression of the full-length dystrophin protein and minimizing the occurrence of revertant fibers, without the loss of other dystrophin isoforms. This was achieved by CRISPR/Cas9-mediated cleavage of the rat *Dmd* within intronic regions that flank exons 22 to 26, resulting in the deletion of approximately 18kb of the sequence within *Dmd* (**Figure 1A**). This deletion, confirmed by PCR (**Figure S1A**), removes 800 nucleotides of protein coding sequence resulting in a premature termination codon in exon 27 (**Figure 1B**). The mutant *Dmd* transcript is predicted to produce a 106 kDa N-terminal dystrophin protein fragment in place of the full-length dystrophin protein. Immunoblotting of muscle lysates with antibodies that recognize the N-terminus of dystrophin demonstrate loss of full-length dystrophin protein in MDR samples (**Figure 1C**) and presence of a truncated N-terminal fragment of the predicted molecular weight (∼106 kDa). Immunofluorescence (IF) of muscle transverse sections also demonstrates the lack of sarcolemmal dystrophin staining in MDR tissues (**Figure 1D**) except in skeletal muscle revertant fibers (**Figure S1B**). The lack of full-length dystrophin protein led to an expected reduction in sarcolemmal localization and protein levels of dystrophin glycoprotein complex (DGC) components– β-dystroglycan, α-dystrobrevin, α-sarcoglycan, and syntrophins (**Figure 1E-F, S1B**). Upregulation of utrophin was also observed (**Figure S1C**). These results confirm the generation of the MDR genetic homolog of DMD that exhibits the expected reduction of sarcolemma-stabilizing DGC complex formation.

**Figure 1:**
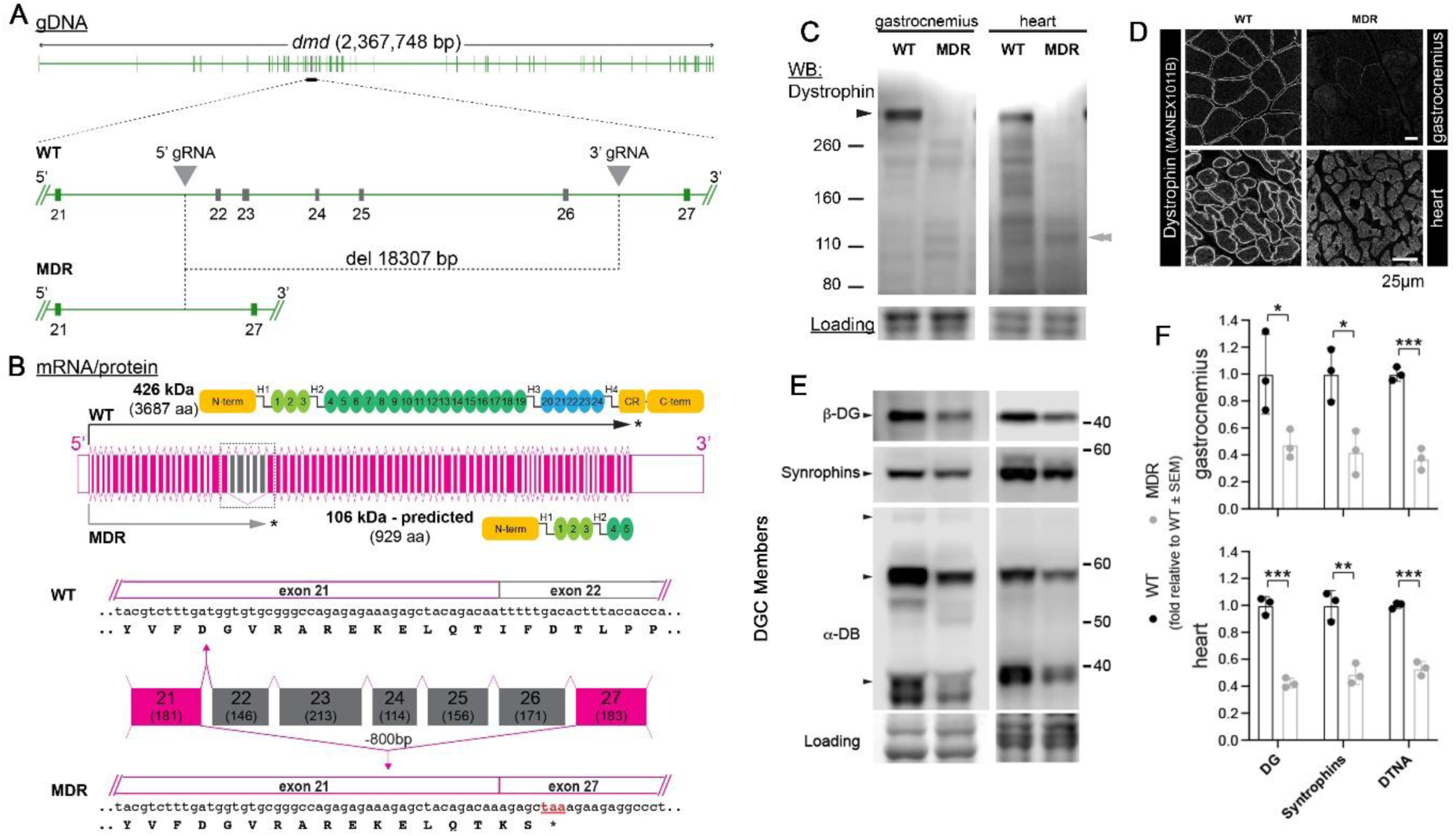
CRISPR/Cas9-mediated deletion of full-length rat dystrophin. (**A**) Organization of the rat *Dmd* that spans ∼2.3 mbp of the X chromosome. The 5 exons targeted for removal – 22-26 (in gray) and the two flanking exons (21 & 27 – in green) are shown in more detail (WT). The two guide RNAs (5’ and 3’ gRNAs) aided in the Cas9-mediated deletion of 18,307 bps that include exons 22-26 (MDR). (**B**) The 5’ and 3’ untranslated regions (unfilled magenta boxes) flank the 79 exons (filled magenta boxes) that contribute coding sequence of full-length dystrophin protein. Indicated with a closer view of exons 21-27 is the number of nucleotides each exon contributes (in parentheses). The 5 deleted exons (in gray) together contribute 800 bp of protein coding sequence, whose removal leads to a frame shift and a premature termination codon within the region of the transcript coded by exon 27 (*, MDR). (**C**) Immunoblotting confirms loss of full-length dystrophin (427 kDa; black arrow) and presence of predicted 106 kDa N-terminal fragment (gray double arrow) in MDR gastrocnemius and heart lysates, and (**D**) immunofluorescence confirms loss of dystrophin staining at the sarcolemma of muscle fibers and cardiomyocytes (scale bars = 25 µm). (**E-F**) Immunoblotting was also performed for dystrophin glycoprotein complex (DGC)members β-dystroglycan (β-DG), α-dystrobrevin (α-DB), and syntrophins. Data are displayed as mean±SEM with individual values and were analyzed using Student’s T-tests (α = 0.05; * p < 0.05, ** p < 0.01, *** p < 0.001).

### Skeletal muscle functional deficits exhibited by the MDR model of DMD

To investigate the functional deficits associated with this rat model of DMD, we assessed WT and MDR *in situ* extensor digitorum longus (EDL) and soleus muscle force production and *ex vivo* diaphragm function at 3, 6, and 12 months of age (mo). As anticipated, significant deficits were found for MDR EDL maximum force production (Po), mass-normalized force production, and resistance against eccentric contraction-induced force drop (**Figure 2A**). While MDR soleus muscles demonstrated significant drops in Po at 3 and 12 mo, mass-normalized force was only significantly lower at 12 mo (**Figure 2A**). *Ex vivo* examination of diaphragm function revealed drastically lower specific tension at all three ages, whereas eccentric contraction-induced force drops did not show significance at these ages (**Figure 2B**). In agreement with these functional deficits associated with the MDR phenotype, rat bodyweight and limb muscle masses were significantly decreased compared to WT controls (**Table 1**), yet heart masses, liver masses, and tibia lengths were not different between the genotypes. Muscle fiber size analysis of the diaphragm and soleus also demonstrate smaller fibers in MDR muscles (**Figure S3**). These data indicate that MDR muscles exhibit considerable functional and morphological deficits compared to WT controls, therefore, MDR represents a large rodent DMD model with a dynamic range that can effectively evaluate potential therapeutics.

**Figure 2:**
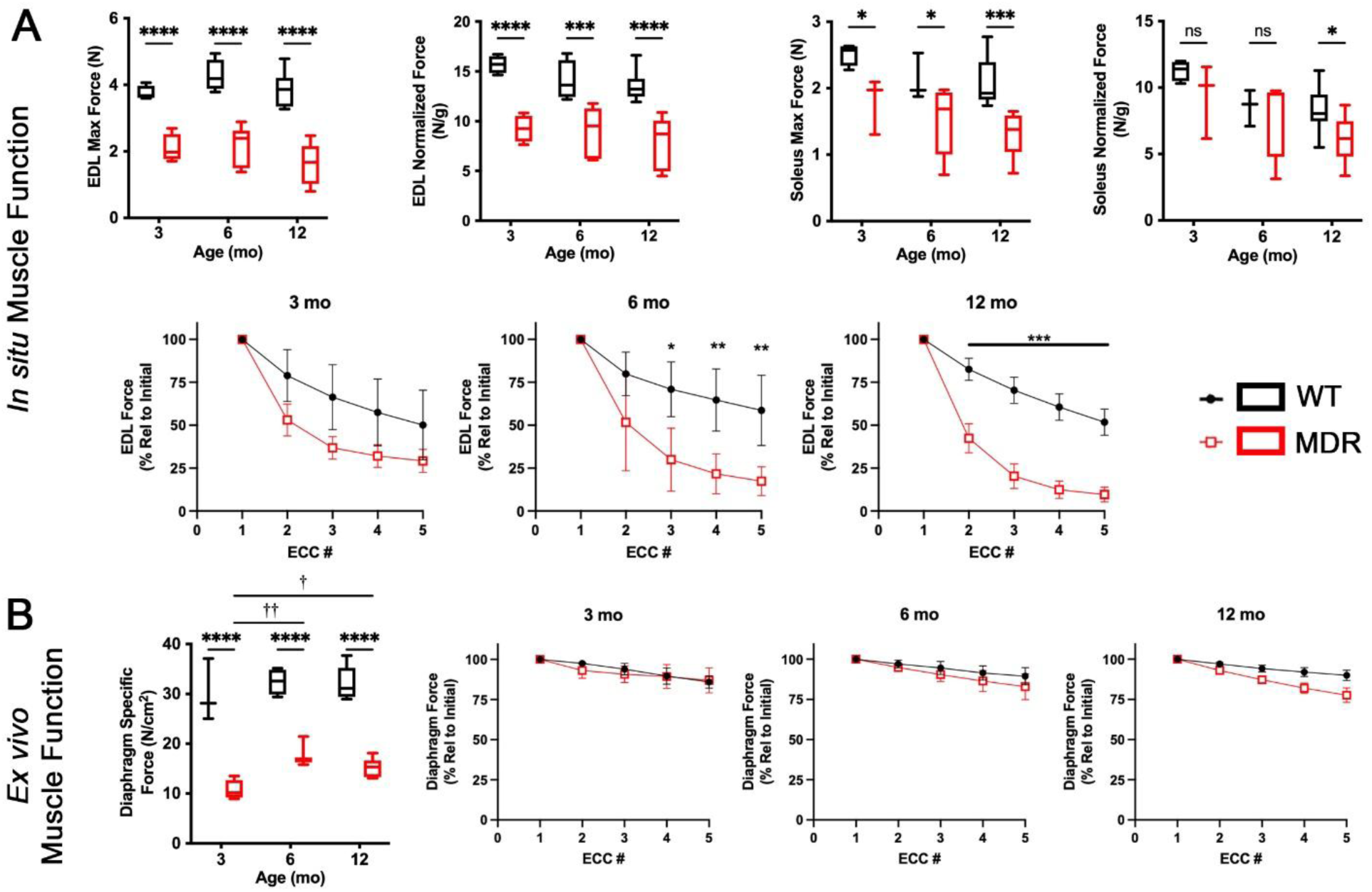
Skeletal muscle function in the MDR model of DMD. (**A**) *In situ* muscle function of the extensor digitorum longus (EDL) shows early and persistent deficits in absolute and normalized force production in muscular dystrophy rats (MDR) at 3, 6, and 12 months of age (mo). Absolute force in soleus was significant for age with lower absolute forces in soleus muscles from MDR at all ages, and significantly lower normalized forces at 12 months of age. Eccentric contractions caused a greater loss of force in MDR EDLs at all ages. (**B**) Isolated diaphragm force production from the same animals was also consistently lower in MDR with significant dependence of age. Loss of force in MDR diaphragms following eccentric contractions was most evident at 6 months of age. Isometric force data are displayed as box-and-whisker plots showing minimum and maximum values and analyzed by 2-way ANOVA followed by Tukey’s post-hoc testing. * p < 0.05, ** p < 0.01, *** p < 0.001 between strains; † p < 0.05, †† p < 0.01 between ages. Eccentric contraction data are mean±SEM, analyzed by multiple T-tests (* p < 0.05, ** p < 0.01, *** p < 0.001).

**Table 1.** Dystrophic rats exhibit reduced body weights and muscle masses. Data are displayed as mean±SEM and were analyzed using 2-way ANOVA for age (†) and strain (*) followed by Tukey’s post-hoc comparisons. (α = 0.05; *,† p < 0.05, **,†† p < 0.01, ***,††† p < 0.001).

| Age (mo) | 3 |  | 6 |  | 12 |  |
| --- | --- | --- | --- | --- | --- | --- |
| Genotype | WT | MDR | WT | MDR | WT | MDR |
| Body weight (g) | 450.0±11.3 | 413.8±9.1* | 520.3±23.7†† | 457.2±13.6***,† | 619.5±21.5††† | 502.8±13.3***,†† |
| EDL Mass (mg) | 236.6±8.0 | 217.9±6.4 | 290.2±9.1†† | 239.2±7.7*** | 272.0±8.0†† | 204.4±10.8*** |
| Soleus Mass (mg) | 212.5±8.6 | 191.0±4.7 | 244.3±9.5 | 202.0±8.7** | 252.9±7.1† | 180.9±11.9*** |
| Tibialis Anterior Mass (mg) | 909.6±30.4 | 856.4±23.1 | 1103.5±17.5††† | 980.2±23.9**,† | 1034.4±23.4† | 849.6±35.8*** |
| Plantaris Mass (mg) | 491.2±19.6 | 442.1±20.2 | 599.0±13.6†† | 488.3±14.2*** | 598.6±15.6††† | 409.4±20.3*** |
| Medial Gastroc Mass (mg) | 1196.4±72.4 | 1090.4±30.1 | 1434.2±70.8 | 1260.5±50.9 | 1316.0±38.7 | 1035.9±70.6*** |
| Lateral Gastroc Mass (mg) | 1393.9±42.4 | 1195.8±64.3* | 1652.9±64.0† | 1346.1±44.4** | 1618.5±49.4† | 1305.8±72.1*** |
| Quadriceps Mass (mg) | 4199.0±152.1 | 3403.8±94.6*** | 5037.6±64.7††† | 3659.6±68.5*** | 4983.9±81.1††† | 3193.8±157.0*** |
| Heart Mass (mg) | 1144.4±58.6 | 1125.6±58.6 | 1498.5±100.4 | 1396.6±89.3 | 1414.9±70.5 | 1360.5±88.1 |
| Liver Mass (mg) | 14279.2±630.7 | 14892.8±636.7 | 15764.8±491.5 | 14598.1±388.2 | 14477.4±479.9 | 15261.4±531.7 |
| Tibia Length (mm) | 44.0±0.5 | 43.1±0.4 | 46.3±0.3 | 46.2±0.2 | 47.7±0.2 | 47.4±0.2 |

**Table 2.** Interspecies comparison of muscle function in 12 month-old preclinical models of DMD.

| DMD Model | Strain (Species) | Muscle/Group | Measure | $\Delta$ from WT (%) | Reference |
| --- | --- | --- | --- | --- | --- |
| MDR | Sprague-Dawley (*Rattus norvegicus*) | Diaphragm | *Ex vivo* SPo | -52.50% | Current Study |
|  |  | EDL | *In situ* Po | -41.60% |  |
|  |  | Soleus | *In situ* Po | -26.10% |  |
| mdx | C57BL/10 (*Mus musculus*) | Diaphragm | *Ex vivo* SPo | -61.80% | Hammers et al. (2020) [15] |
|  |  | EDL | *Ex vivo* Po | +4.60% |  |
| D2.mdx | DBA/2J (*Mus musculus*) | Diaphragm | *Ex vivo* SPo | -52.00% |  |
|  |  | EDL | *Ex vivo* Po | -45.93% |  |
| GRMD | Golden Retriever (*Canis lupus familiaris*) | Distal pelvic limb | *In vivo* extension | -63.97% | Kornegay et al. (1999) [29] |
|  |  |  | *In vivo* flexion | -61.92% |  |

To attempt to explain the lack of pronounced functional deficits exhibited by the MDR soleus, we performed force-stimulation frequency analyses for 12 mo WT and MDR diaphragm, EDL, and soleus muscles. The stimulation frequency for highest force output for all three muscles from WT rats was 75 Hz (**Figure 3A**). For MDR diaphragm and EDL, this remained at 75 Hz. However, for MDR soleus muscles, there was a right shift in the force-frequency curve resulting in an optimal stimulation frequency of 125 Hz. Such shifts may be explained by altered muscle fiber type composition within the affected muscles. In agreement, we observed a clear slow-to-fast fiber type shift in soleus muscle of MDR animals (**Figure 3B-C**). While essentially no type IIx fibers were present in the soleus muscle of WT rats, type IIx fibers accounted for more than 10% of all muscle fibers present in MDR soleus muscles (**Figure 3C**). No such alteration in fiber types was observed with the EDL muscle. Thus, the right shift in the force frequency curve of MDR soleus muscle compared to WT is concurrent with the slow-to-fast shift in MDR soleus muscle fiber types. This may account for the reduced functional deficits identified in the MDR soleus.

**Figure 3:**
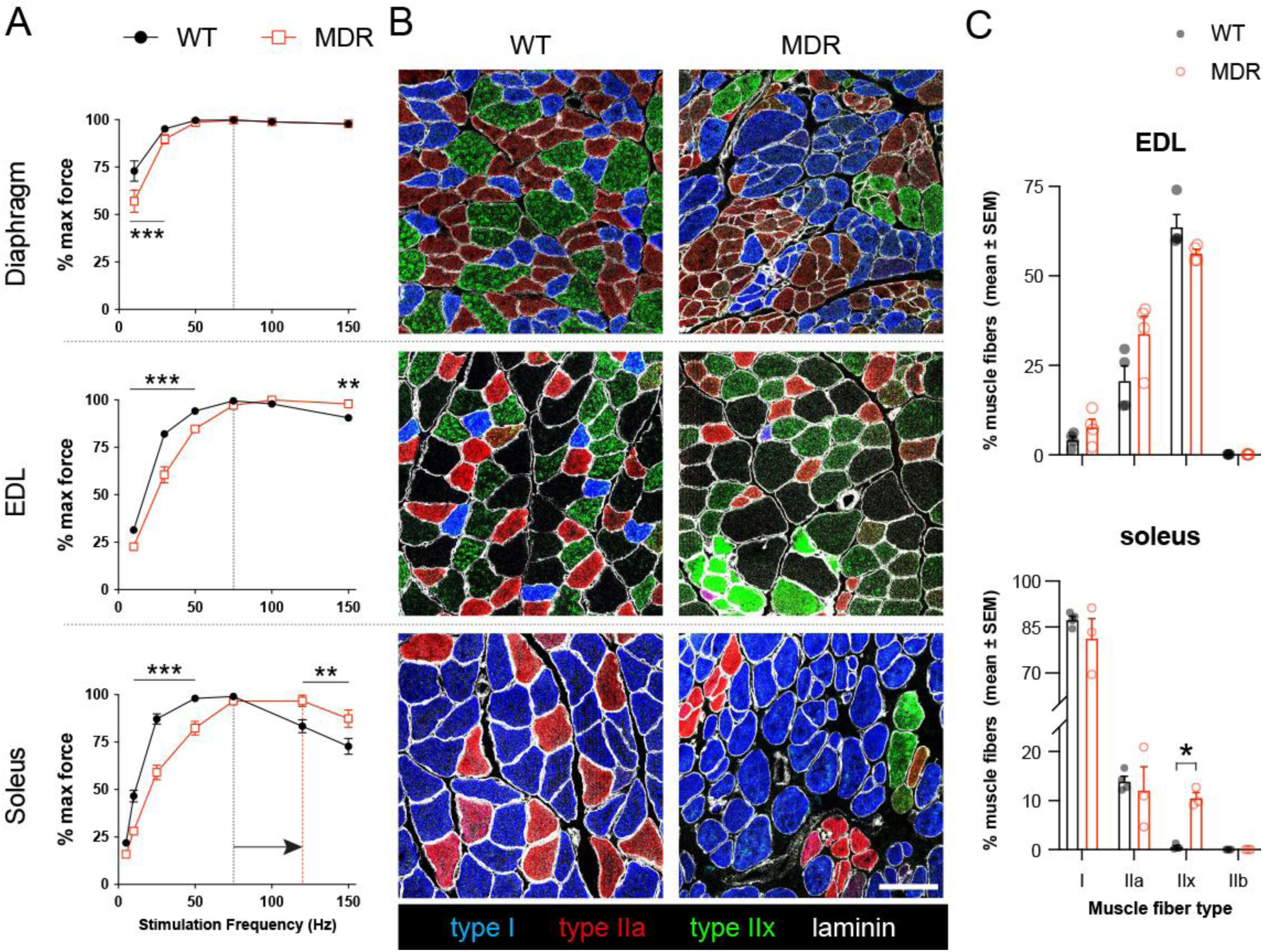
A shift in optimal stimulation frequency of MDR soleus muscle concurrent with an altered fiber type composition. (**A**) The stimulation frequency for the greatest force output was 75 Hz in all wild-type (WT) muscles and dystrophic (MDR) diaphragm and EDL muscles. For MDR soleus muscle, however, there was a right shift in the force-frequency curve and an increase of the optimal stimulating frequency to 125 Hz. (**B,C**) Myosin heavy chain isoform-specific antibody-mediated labeling of muscle transverse sections (**B**) revealed a slow-to-fast shift in the composition of muscle fiber types with the emergence of type IIx fibers in MDR soleus muscles (**C**; Scale bar = 100 µm). No significant changes were observed with MDR EDL myofiber type composition compared to WT. Data are displayed as mean±SEM and were analyzed using Student’s T-tests (α = 0.05; * p < 0.05, ** p < 0.01, *** p < 0.001).

### Histopathologic features of MDR skeletal muscle

In addition to functional measures, we performed histological assessments on WT and MDR muscles to evaluate degeneration/regeneration, fibrosis, and fat deposition. As expected in dystrophic muscles, MDR diaphragm, soleus, and gastrocnemius muscles exhibited muscle fiber disorganization, degeneration, and inflammatory infiltration, as visualized by H&E staining (**Figures 4A and S3A**), as well as pronounced increases in endomysial fibrosis, as seen with PSR staining (**Figures 4B and S3A**). While these features were uniform in the diaphragm and soleus, they were more heterogeneous in the gastrocnemius (**Figure S3A**). Quantification of whole muscle images of PSR stained cryosections revealed significantly more fibrosis in the 3 and 6 mo MDR soleus and diaphragm (**Figure 4C**). Specific pathological markers were also investigated in WT and MDR muscles via IF staining. For instance, positive staining for embryonic myosin heavy chain, a marker of active regeneration, was found in 4 mo diaphragm and soleus sections (**Figure 4D**), confirming regenerative fibers in MDR muscles. Furthermore, MDR soleus muscles exhibit increased staining for the fibroblast marker PDGFRα, the adipocyte marker perilipin (PLIN), the myofibroblast and smooth muscle marker α-smooth muscle actin (α-SMA), and the matricellular proteins periostin, cartilage oligomeric matrix protein (COMP), and SPARC-related modular calcium binding protein 2 (SMOC2; **Figure 4E and S3B-C**). Therefore, MDR muscles display major dystrophic histopathological features in addition to their major functional deficits.

**Figure 4:**
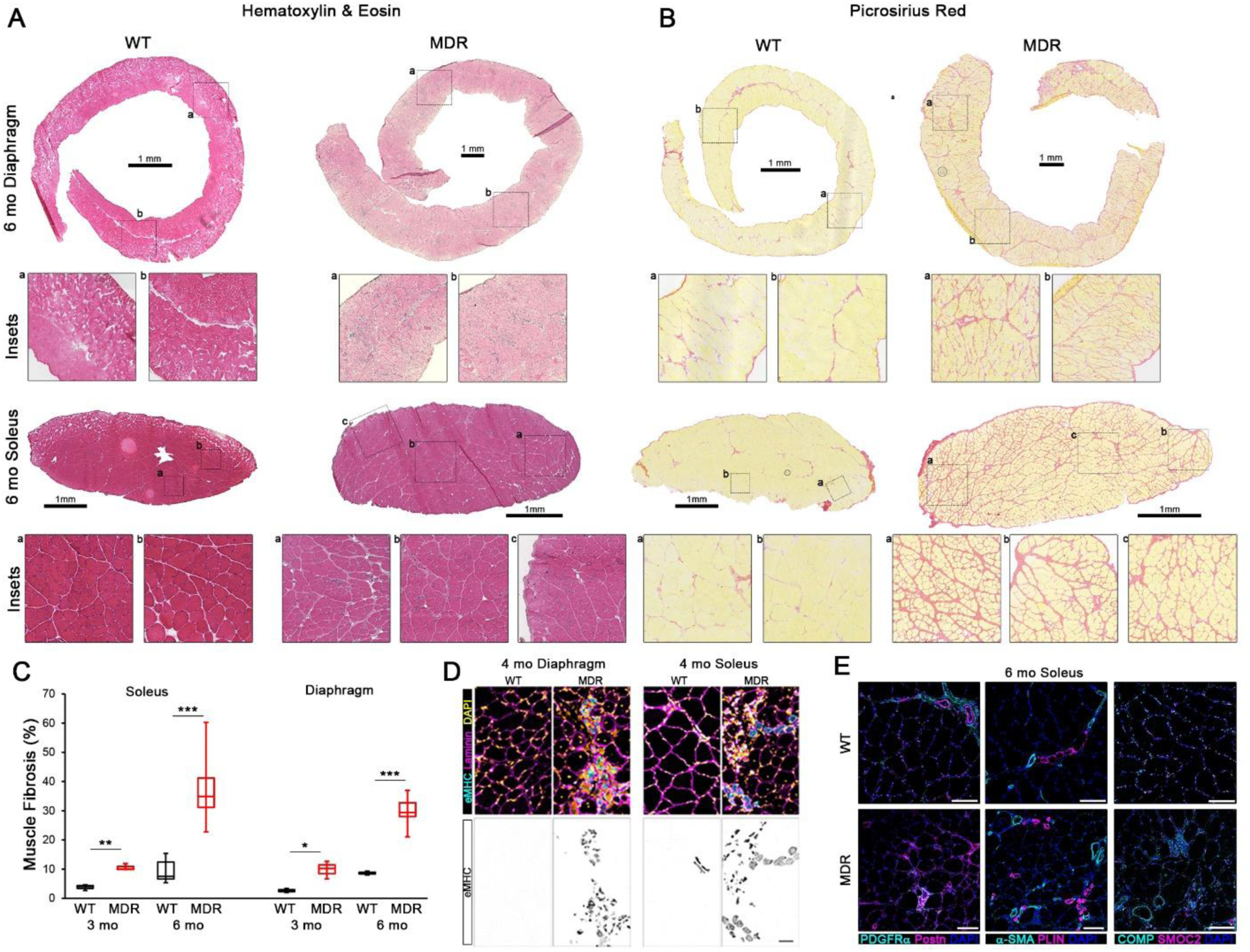
Histopathological features are exhibited by MDR skeletal muscle. Histological assessments were performed on wild-type (WT) and dystrophic (MDR) diaphragms and soleus muscles using (**A**) hematoxylin & eosin and (**B**) picrosirius red staining (PSR; Scale bars = 1 mm). (**C**) Muscle fibrosis was quantified from PSR-stained slides. Immunofluorescence was used to investigate specific histopathological features associated with MDR muscles, including (**D**) embryonic myosin heavy chain (eMHC; scale bar = 50 µm) to identify regenerating muscle fibers in the diaphragm and soleus, as well as (**D**) the fibrosis and adipose associated markers of PDGFRα, periostin (Postn), α-smooth muscle actin (α-SMA), perilipin (PLIN), COMP, and SMOC2 (scale bars = 100µm). Data are displayed as box-and-whisker plots showing minimum and maximum values and were analyzed using Student’s T-tests (α = 0.05; * p < 0.05, ** p < 0.01, *** p < 0.001).

Previous reports of histopathology associated with rat models of DMD suggest that the development of intramuscular fat, an aspect of human pathology not found in other animal models, is a feature of dystrophic rat muscle [20–22]. We also observed modest adipose deposition in MDR soleus muscles (**Figures 4A, 4E, S3B-C**), motivating the investigation of adiposity across MDR muscles. In addition to PLIN positive staining interspersed between soleus muscle fibers (**Figure S4A**), immunoblotting revealed a significant increase in PLIN content in 12 mo MDR gastrocnemius muscles, compared to WT (**Figure S4B**). While there was no MDR-associated PLIN staining observed in 3 and 12 mo diaphragms, increased adipogenesis was observed in MDR EDL and soleus muscles at these ages (**Figure S4C**). However, PLIN^+^ staining accounted for less than 1% of total muscle area, even in the most extreme instances. Therefore, any increase in intramuscular fat we observed in the MDR muscles appears unlikely to meaningfully impact the functional output of the diseased muscles in MDR and does not model the situation in human DMD muscles.

## DISCUSSION

Effective development of therapeutics for human disease requires preclinical models that **a**) accurately portray major pathologies found in the clinic and **b**) are feasible to implement in rigorous studies to test efficacy using clinically relevant outcomes. The goal of the present study was to thoroughly investigate the functional skeletal muscle pathology associated with a CRISPR/Cas9 generated rat model of DMD. We identify significant deficits in muscle mass and function associated with MDR skeletal muscles, particularly the diaphragm and EDL, thus this model offers a dynamic range suitable for therapeutic evaluation by 3 months of age. Furthermore, MDR muscles exhibit progressive histopathological development without an overt degenerative phase early in life. These findings indicate the MDR model of DMD is an intermediately-sized preclinical model to investigate DMD therapeutic development using rigorous functional outcomes.

Preclinical animal models of human genetic disease have been the mainstay of understanding mechanisms of disease and leading to development of cures. DMD is no exception, and progress in the field has been aided largely by mouse models. The *mdx* mouse is a genetic homolog of the human disease, with loss of dystrophin arising from nonsense mutation [23]. Hence, there is much that has been learned about the basis of DMD pathology through the more than 3800 published studies based on the *mdx* mouse. Even so, the C57BL/10ScSn *mdx* mouse, the most prevalent animal model of DMD, fails to reliably replicate several key aspects of human DMD pathophysiology, including sustained atrophy and force producing capacity of limb muscles (**Table 4**), ambulatory deficits, presentation of cardiomyopathy as well as reduced life-expectancy [9–12, 15]. These aspects have limited the translational value of this model of DMD.

Strategies to increase disease severity in mouse models include use of the DBA/2J background (D2.*mdx*), resulting in weaker and more fibrotic muscles that better represent DMD muscle disease. However, there is still an early wave of muscle degeneration that initiates the onset of muscle pathology in D2.*mdx* mice [15, 24]. Although the mouse model lacking both dystrophin and utrophin (*mdx*;*utrn*^-/-^) produces a more severe pathophysiology with shorter life-spans [25, 26], they are not a genetic homolog of the human disease, precluding them from potential therapeutic strategies that involve upregulation of endogenous utrophin. Furthermore, these mouse models exhibit minimal adipose infiltration, which is a prominent feature of DMD muscle.

Canine models, including the golden retriever muscular dystrophy (GRMD) model [27–29], present with a more severe disease progression in part due to their larger size. However, considerable phenotypic variation between individual GRMD animals [19, 30–33], as well as the significant care burden for these severe models make them cost prohibitive for general use in preclinical studies. Smaller animals used to model DMD, such as *C. elegans*, drosophila and zebrafish, while useful for high-throughput screening of compounds *in vivo* [34], have limited utility in modeling human pathophysiology. Overall, organism scale is an important consideration in deciding which preclinical model is optimal for a particular study.

Due to these considerations, recent efforts have modeled dystrophin deficiency in rats [20–22]. Each reported inactivation of rat *Dmd* targeted a distinct region of the gene – exon(s) 3/16 [21], 23 [22] or 52 [20] – that led to the loss of FL dystrophin expression and reproduced some aspects of the human DMD pathology. All three DMD rat models show increased fibrosis of striated muscle tissues, evidence of intramuscular fat, and deficits in motor tasks that suggest reduced force generating capacity of skeletal muscles. Additionally, inactivation of *Dmd* exon 23 or 52 results in reduced body weight and muscle mass. These studies support the use of the rat models for examining the pathophysiological mechanisms of the disease and potential therapeutic strategies: a cost-effective and genetically homogeneous rodent model with a severe dystrophic phenotype that mimics many aspects of human pathophysiology.

The current study substantiates that the MDR model shares the dystrophic hallmarks found in other rat models of DMD, including loss of dystrophin, reduced DGC content, and progressive histopathology. Importantly, we advance the understanding of rat DMD models by confirming that MDR muscles exhibit considerable functional deficits in muscle contractile force production across the lifespan of the rat, which is comparable to the more severe models mentioned above (D2.*mdx* and GRMD, **Table 4**). The initial reports [20–22] on the dystrophin-deficient rat model suggested that there could be fatty replacement of muscles, like the situation in human DMD. However, our study demonstrates that there is minor fatty deposition within the muscles. Thus, like the mouse and dog models of DMD, progressive fibrosis occurs but fatty replacement of muscles does not, unlike in the human disease. Nonetheless, therapeutic testing using this model can utilize this considerable dynamic range of functional deficits to better evaluate potential translational functional benefits that will facilitate informed clinical trial design. Collectively, these findings indicate that the MDR, and likely other rat models of DMD, exhibits adequate skeletal muscle functional and structural decrements to have high utility for investigating therapeutic impact of potential DMD treatments.

## MATERIALS & METHODS

### Animals

All animal studies were approved and conducted in accordance with the University of Florida IACUC. CRISPR/Cas9-mediated *Dmd* inactivation to generate the MDR line was performed by Cyagen, which was maintained on the Sprague-Dawley rat strain (Charles River). Intronic sequences present between exons 21 and 22 as well as 26 and 27 of the rat *Dmd* were specifically targeted for Cas9-mediated cleavage by the following gRNAs (protospacer adjacent motif sequence underlined): 5’-GTCTAATAGTAGGTGATAAG<u>AGG</u>-3’, 5’-CAGCTCTTGTACCCGATTGCTGG-3’. Five potential off-target sites were identified (mismatched nucleotides are underlined) – 3 for the 5’ gRNA (GTCT<u>TG</u>TAGTAGGTGA<u>C</u>AAGTGG on chromosome 7, GT<u>G</u>T<u>G</u>ATAGTAGGTGA<u>C</u>AAG GGG on chromosome 2, GTCTAA<u>G</u>AG<u>C</u>AGG<u>G</u>GATAAGTGG on chromosome 17) and 2 for the 3’ gRNA (<u>AG</u>GCTCTTGT<u>C</u>CCCGATTGCAGG on chromosome 16, CAGCTCTTGT<u>G</u>CC<u>G</u>GA<u>A</u>T GCAGG on chromosome 5). The results of PCR and sequence analysis indicated that no off-targets indels were produced at these sites. Rats were housed up to two littermates per cage, randomly assigned into pairs, provided *ad libitum* access to food (NIH-31 formulation diet), water, and enrichment, and maintained on a 12-hour light/dark system.

### MDR genotyping

DNA was isolated from rat tail snips using DNeasy Blood and Tissue Kit (Qiagen; Cat #69506). Genotypes were confirmed by standard polymerase chain reaction (30 amplification cycles with an annealing temperature of 60°C) using the following primer sets: Dmd-F 5’-GGGATTCACTGA AGGTACAAAATCTACA-3’; Dmd-WT-R 5’-GTAGTATGCTTAATCACCAACCCCAG-3’; Dmd-MDR-R 5’-GGGTGAGGAAGACTTGTAATAGTGTCTTC-3’. Presence of a WT allele results in a 586 bp product (Dmd-F & Dmd-WT-R), and a mutant allele in a 790 bp product (Dmd-F & Dmd-MDR-R).

### In situ & ex vivo muscle function

Rats were anesthetized using 3-3.5% isoflurane and maintained at 2.5-3% for the duration of *in situ* muscle function tests of soleus and EDL muscles. The body temperature was maintained at 37°C with the use of a heating pad and a temperature-controlled animal stage. Throughout the procedure, warm mineral oil and/or Ringer’s solution was applied to the incision sites to protect the tissue from dehydration. An incision was made along the biceps femoris muscle to expose the sciatic nerve, which was cut to allow stimulation of the soleus and EDL muscles through the distal length of the severed sciatic nerve while preventing back-propagation. Muscles were stimulated using a hook electrode. The soleus and EDL were isolated free from other muscles and a suture was secured at the myotendinous junction. On the temperature-controlled animal stage, the knee was fixed to the platform with a screw through an opening made under the patellar tendon and the hind paw secured at a right angle to the platform with medical tape. The loop placed at the myotendinous junction of the isolated EDL or soleus muscle was secured to the lever arm of the tension transducer with the resting tension of the muscle (305 muscle lever system; Aurora Scientific). Following *in situ* measurements, *ex vivo* muscle mechanics were performed on the diaphragm, as previously described [15, 17, 18].

For each of the above preparations, the optimum length (L_O_) of the muscle was found by performing a series of twitches, each with an incremental increase in the length of the muscle, until the length that yielded the maximum twitch force was found. Subsequently, each muscle preparation was subjected to a maximum isometric force test, force-frequency test, and/or a series of eccentric contractions (ECCs). The isometric force tests were performed by stimulating the EDL at 120Hz and the soleus at 100Hz, both for 500ms, with a 180 second rest period between stimulations. The force-frequency test was performed using 500 ms trains with stimulation frequencies ranging from 5 to 150Hz for 500ms each with a resting period of 180 seconds between stimulations. P_O_ is not reported for diaphragm as the preparation involves arbitrary incisions to generate diaphragm strips. Thus, the specific tension (P_O_ normalized by muscle cross sectional area) is the only measure of force production reported for the diaphragm muscle. Eccentric contractile stimulations (ECC) are performed by subjecting the muscle to an isometric contraction at 80hz, with a stretch protocol of 10% of the L_O_ during the last 200ms of contraction. There is a recovery period of 300 seconds between contractions. Values are measured from isometric contraction to isometric contraction, not total force produced during the stretch portion of the stimulation.

### Immunoblotting

Protein extraction and immunoblotting were performed on snap-frozen heart and gastrocnemius muscles, as previously described [16–18], using the following primary antibodies: dystrophin [1:100; MANHINGE1B, Clone 10F9; Developmental Studies Hybridoma Bank (DSHB)], dystrophin (1:100; MANEX1011B; Clone 1C7; DSHB), dystrophin (1:100; MANEX1011C; Clone 4F9; DSHB), β-dystroglycan (1:1000; 11017-1-AP; ProteinTech), dystrobrevin (1:1000; #610766; BD Biosciences), syntrophins (1:2000; #11425; Abcam), and perilipin 1 (1:500; #9349; Cell Signaling Technology). Membranes were stained with Ponceau S to control for equal protein loading and for normalization. The Band signal intensities were measured using Image Studio Lite software (Li-Cor Biosciences), normalized to sample loading (Ponceau S stain), and reported relative to respective control samples.

### Immunofluorescence and histological evaluations

IF was performed as previously described [18] using fresh-frozen OCT-embedded diaphragm, extensor digitorum longus, gastrocnemius and soleus muscles cross-sectioned at 10 µm and fixed in either ice-cold acetone or 4°C 4% paraformaldehyde. The following primary antibodies were used for immunofluorescence in the present study: dystrophin (1:100; MANEX1011B, Clone 1C7; DSHB), β-dystroglycan (1:100; MANDAG2, Clone 7A11; DSHB), utrophin (1:50; VP-U579; Vector Laboratories), α-sarcoglycan (1:50; VP-A105; Vector Laboratories); syntrophins (1:2000; #11425; Abcam), perilipin 1 (1:500; #9349; Cell Signaling Technology), anti-Laminin β2 antibody (1:10,000; [35]), embryonic MHC (1:100; F1.652; DSHB), type I myofibers (1:50; BA-D5; DSHB), Type IIa myofibers (1:50; SC-71; DSHB), type IIx myofibers (1:20; 6H1; DSHB), pan-muscle MHC(1:100; MF 20; DSHB), COMP (1:800; #NBP2-92658; Noveus Biologicals), α-SMA (1:1000; #7817; Abcam), SMOC2 (1:500; #AF5140; R&D Systems), and PDGFRα (1:500; #AF1062; R&D Systems). Image acquisition was performed with a Leica Application Suite X software on a Leica STELLARIS 5 confocal system. Comparative images were stained, imaged, and processed simultaneously under identical conditions. Muscle fiber minimum Feret’s diameter measurements from fluorescence images were performed with a combination of Cellpose segmentation algorithm [36] and LabelsToRois plugin for FIJI/ImageJ [37]. The percent frequency distribution of muscle fiber diameters was plotted in steps of non-cumulative 10-µm bins and were then each fitted with a best-fit curve through a non-linear regression of the raw data.

Hematoxylin & Eosin (H&E) and Picrosirius Red (PSR) staining of muscle sections was performed as previously described [15, 16]. were visualized with a Leica DMR microscope, and images were acquired using a Leica DFC310FX camera interfaced with Leica LAS X software, as well as the Keyence X-BZ All-In-One Microscope (Keyence Corporation, Osaka, Japan). Fibrosis was quantified using Hue and Saturation masks in ImageJ (NIH).

### Statistics

Data were analyzed using two-tailed Student’s T-tests (α = 0.05) or two-way ANOVA (Tukey post-hoc tests; α = 0.05), where appropriate, and are displayed as or min-to-max box-and-whisker plots with the 2^nd^ and the 3^rd^ quartiles within the box along with a line indicating the median, bar graphs or line plots with mean ± standard error of means (SEM), or as histograms. A P value less than 0.05 was considered significant.

## ACKOWLEDGMENTS

We thank Grace Gloger, Kara Pugliese, and Christina Volpe for their technical assistance, and Dr. Alan Russell of Edgewise Therapeutics for generously sharing the dystrophin-deficient rats described in this study. This work was supported by a NIH/NIAMS Wellstone Muscular Dystrophy Specialized Research Center grant (P50-AR052646) to HLS, DWH, and ERB, and a Parent Project Muscular Dystrophy grant to HLS.

## SUPPLEMENTARY FIGURES

**Figure S1.**
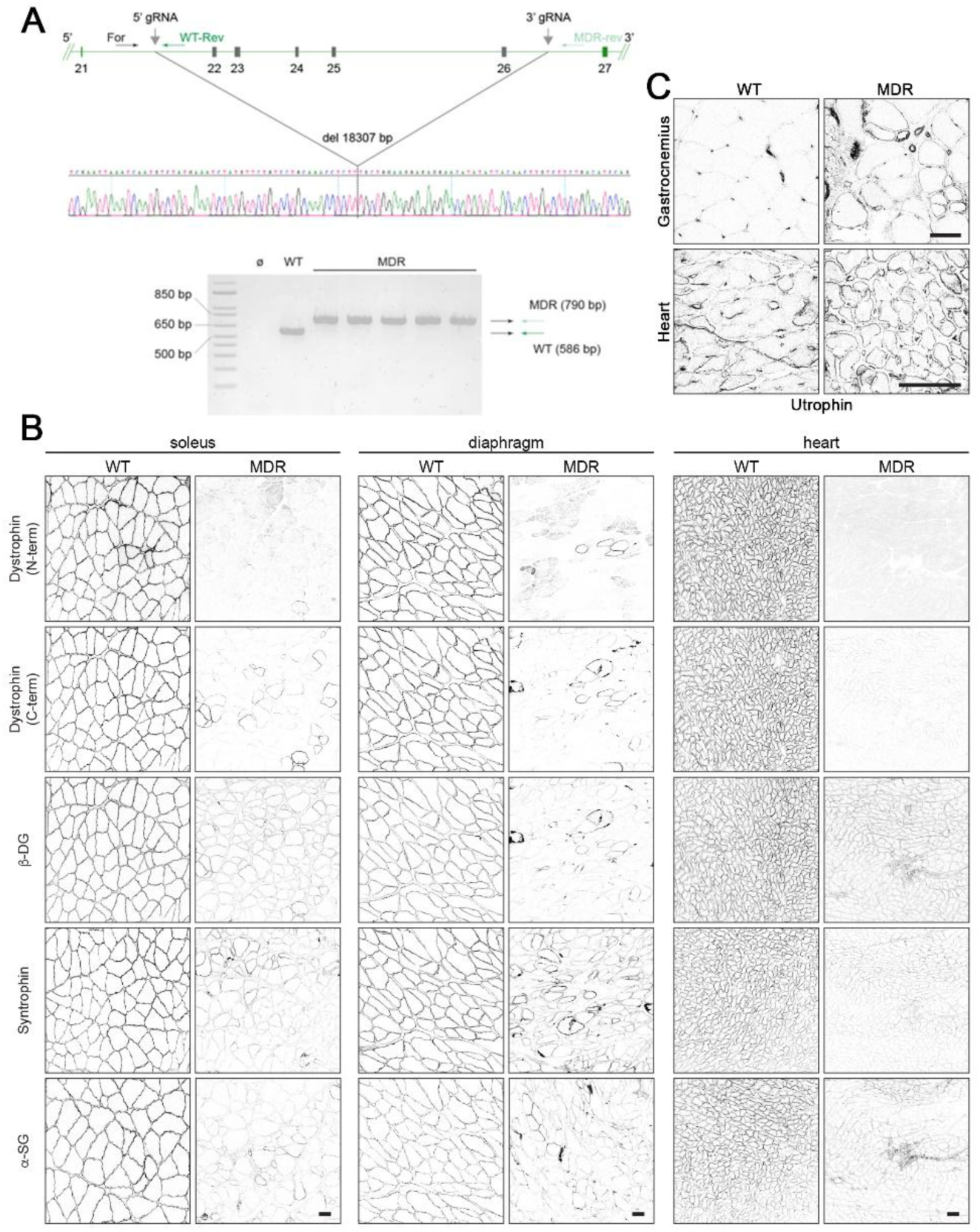
MDR genotyping and DGC localization. (**A**) A set of three primers are used to genotype the rats. For WT, two primers placed within the intronic region between exons 21 and 22a common forward primer (For) and a WT reverse primer (WT rev) situated 5’ and 3’, respectively, to the sequence targeted by 5’ gRNA yield a 568 bp product. For MDR, For and a MDR reverse primer (MDR rev) – latter situated 3’ to the 3’ gRNA – yield a 790 bp product. Examples of the PCR products that match predicted sizes are visible after separation within an agarose gel. A portion of the sequence trace of the 790 bp products confirms the deletion of 18,307-bp stretch of rat *Dmd* that includes exons 22-26. (**B**) Immunofluorescence was used to investigate sarcolemmal localization of dystrophin and the DGC members of β-dystroglycan (β-DG), α-dystrobrevin (α-DB), α-sarcoglycan (α-SG), and syntrophins in WT and MDR soleus, diaphragm, and heart, as well as (**C**) utrophin localization in the WT and MDR gastrocnemius and heart (scale bars = 50 µm).

**Figure S2.**
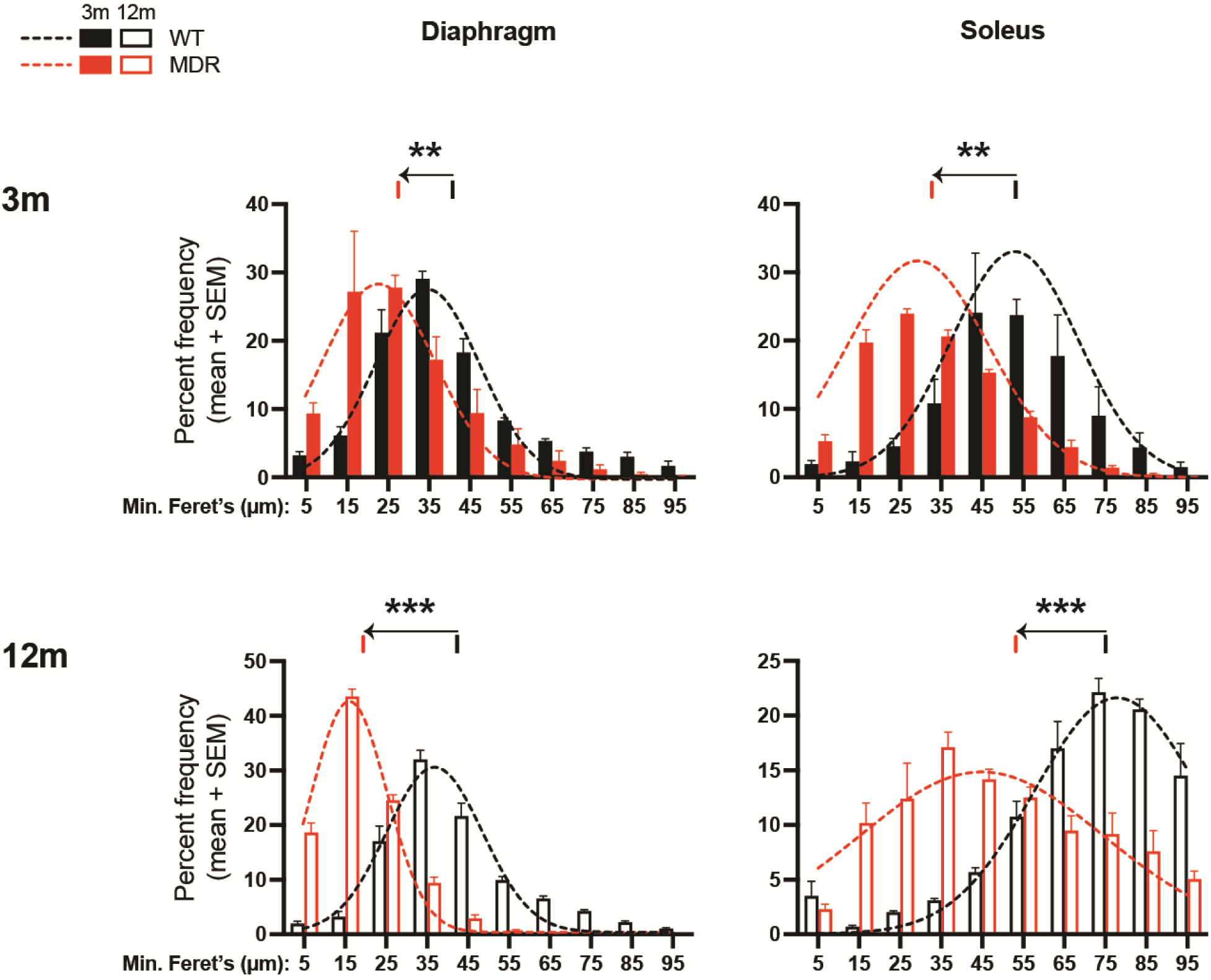
Reduced myofiber diameters of MDR skeletal muscles. The frequency distribution of WT and MDR diaphragm and soleus muscle fibers minimum Feret’s diameter was performed at 3 and 12 months of age. There is a left shift (towards smaller diameter) for dystrophic rat muscle fibers. The vertical bars above the frequency distribution plot indicate the average minimum Feret’s diameter, with the arrow indicating the WT-to-MDR shift. Data are displayed as mean±SEM and were analyzed using multiple T-tests (α = 0.05; * p < 0.05, ** p < 0.01, *** p < 0.001).

**Figure S3.**
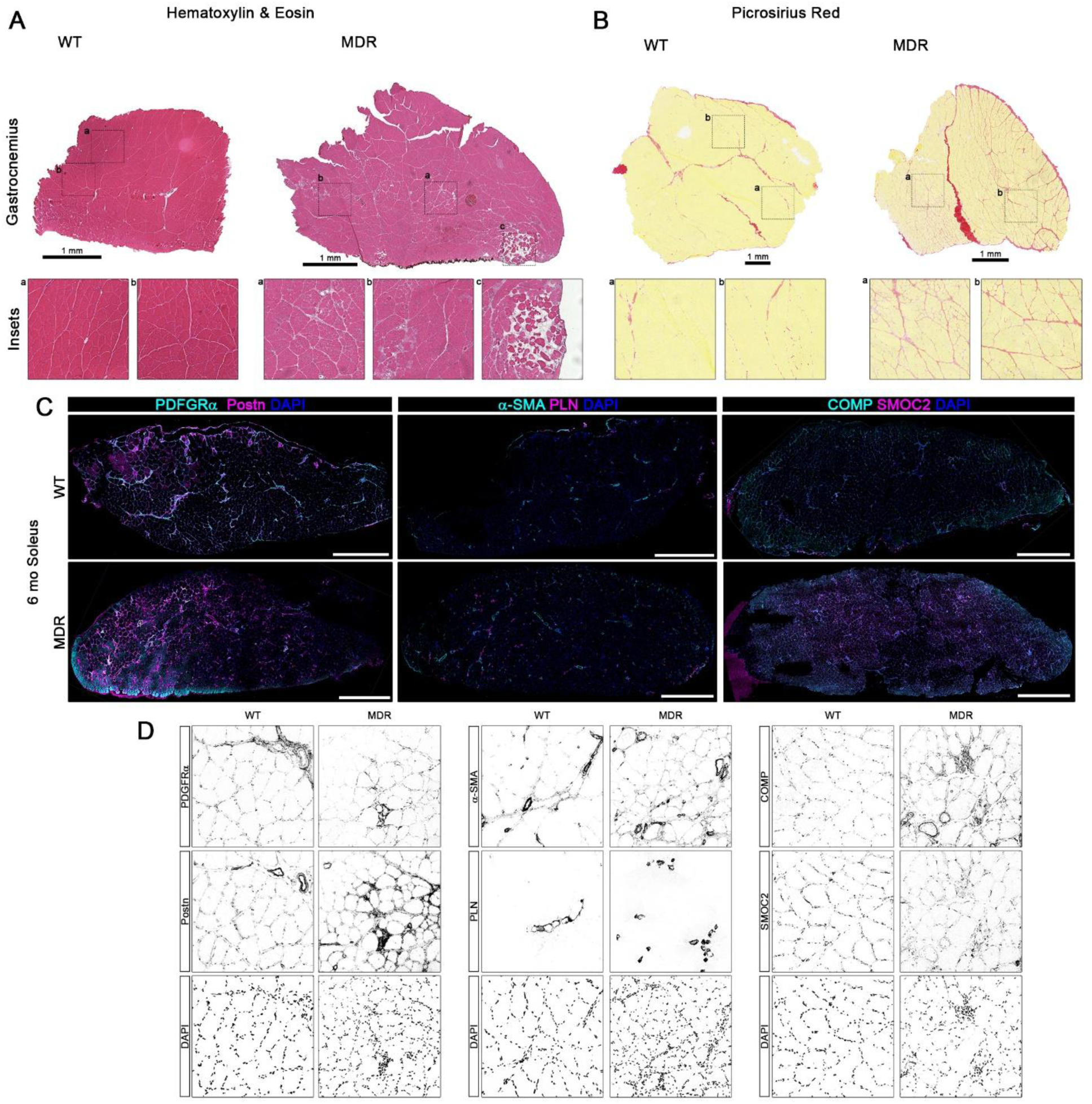
Histopathological features in MDR muscle. Histology was performed on 6 month-old wild-type (WT) and dystrophic (MDR) gastrocnemius muscles using (**A**) hematoxylin & eosin and (**B**) picrosirius red (scale bar = 1 mm). (**C-D**) Immunofluorescence was performed on 6 month-old soleus muscles to investigate the localization of PDGFRα, Postn, α-SMA, PLIN, COMP, and SMOC2 (scale bars = 1 mm). The merged images of (**D**) are located in **Figure 4E**.

**Figure S4:**
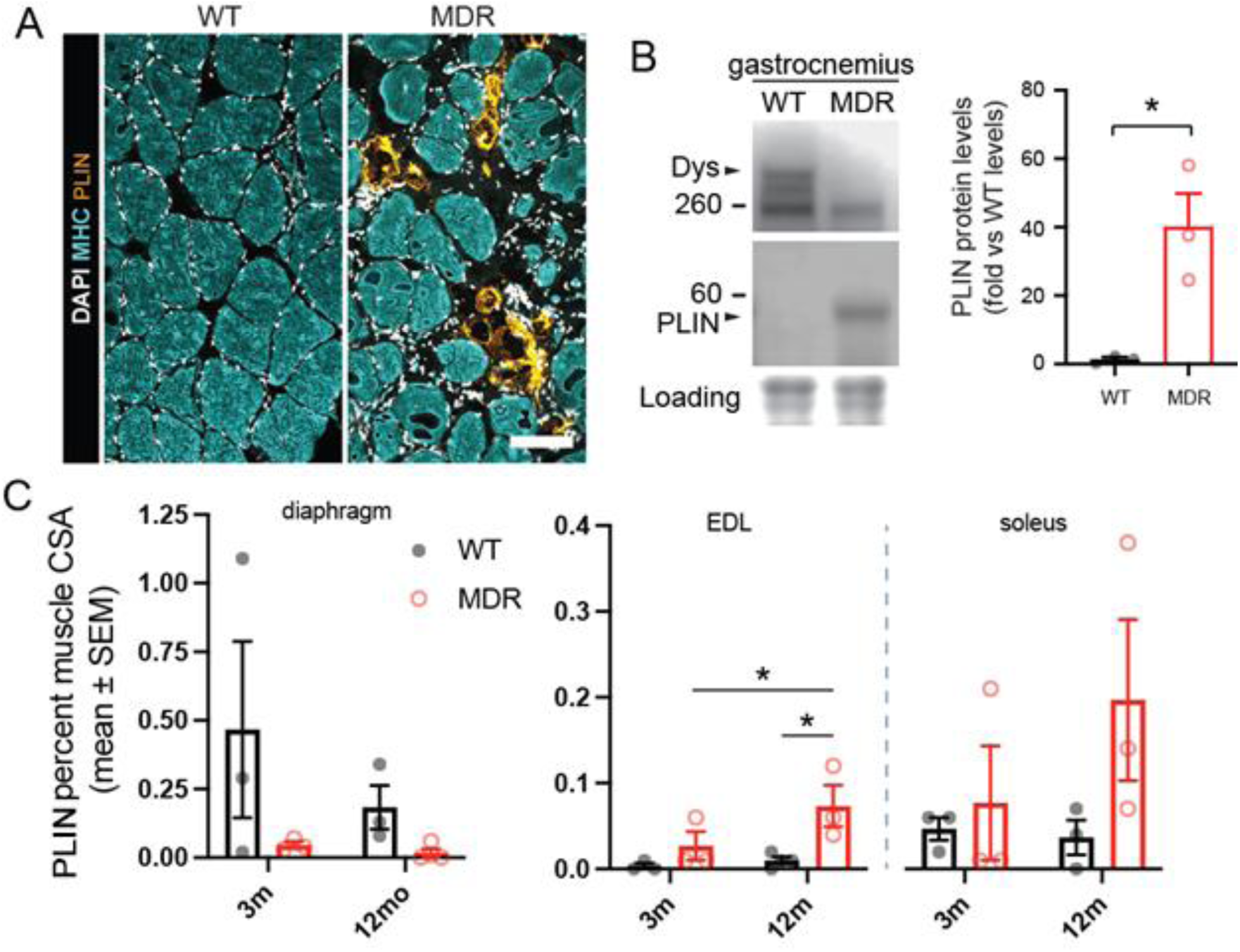
Adipocytes in MDR skeletal muscles. (A) Transverse sections of the soleus muscle labeled for myosin heavy chain (MHC) and perilipin 1 (PLIN) reveal the presence of intramuscular fat deposition in the dystrophic animals (Scale bar = 100 µm). (B) Western blots of gastrocnemius muscle lysates also show increased PLIN protein expression with MDR vs WT rats by Student’s T-test. (C) Analysis of muscle transverse sections stained for perilipin shows an increase in the PLIN-positive fraction of muscle cross sectional area in 12-month-old MDR EDL muscles, but not in diaphragm or soleus, by 2-way ANOVA for age and strain following by Fisher’s LSD test. Data are displayed as mean±SEM with individual values.

## Notes

### Competing Interest Statement

The authors have declared no competing interest.

